# Glycocalyx engineering with heparan sulfate mimetics attenuates Wnt activity during adipogenesis to promote glucose uptake and metabolism

**DOI:** 10.1101/2021.07.08.451710

**Authors:** Greg W. Trieger, Ariane R. Pessentheiner, Sean C. Purcell, Courtney R. Green, Natalie DeForest, Karl Willert, Amit R. Majithia, Christian M. Metallo, Kamil Godula, Philip L. S. M. Gordts

**Affiliations:** Department of Chemistry and Biochemistry, University of California, San Diego, La Jolla, CA 92093, USA; Department of Medicine, Division of Endocrinology and Metabolism, University of California, San Diego, La Jolla, CA 92093, USA; Department of Bioengineering, University of California, San Diego, La Jolla, CA 92093, USA; Department of Cellular and Molecular Medicine, University of California, San Diego, La Jolla, CA 92093, USA; Department of Pediatrics, University of California, San Diego, La Jolla, CA 92093, USA; Glycobiology Research and Training Center, University of California, San Diego, La Jolla, CA 92093, USA

**Author notes:** **Corresponding authors**: Kamil Godula, Department of Chemistry and Biochemistry, University of California San Diego, 9500 Gilman Drive #0358, La Jolla, CA 92093-0358, Philip L.S.M. Gordts, Department of Medicine, University of California San Diego, 9500 Gilman Drive #0678, La Jolla, CA 92093-0678.

## Abstract

Adipose tissue (AT) plays a crucial role in maintaining me tabolic homeostasis by storing lipids and glucose from circulation as intracellular fat. As peripheral tissues like AT become insulin resistant, decompensation of blood glucose levels occurs causing type 2 diabetes (T2D). Currently, glycocalyx modulating as a pharmacological treatment strategy to improve glucose homeostasis in T2D patients is underexplored. Here, we show a novel role for cell surface heparan sulfate (HS) in establishing glucose uptake capacity and metabolic utilization in differentiated adipocytes. Using a combination of chemical and genetic interventions, we identified that HS modulates this metabolic phenotype by attenuating levels of Wnt signaling during adipogenesis. By engineering the glycocalyx of preadipocytes with exogenous synthetic HS mimetics, we were able to enhance glucose clearance capacity after differentiation through modulation of Wnt ligand availability. These findings establish the cellular glycocalyx as a possible new target for therapeutic intervention in T2D patients by enhancing glucose clearance capacity independent of insulin secretion.

**SIGNIFICANCE:** Metabolic disorders associated with the Western-style diet, such as type 2 diabetes, are among the main drivers of mortality in the US and globally, with more than 380 million people currently affected by this disease worldwide. However, treatment options for type 2 diabetes are currently limited to management of caloric uptake and expenditure, with none able to reverse the condition long-term. The ability to reprogram adipose tissues to improve their overall capacity to clear glucose may provide one such opportunity. Here we provide evidence that glycocalyx remodeling in pre-adipocytes with heparan sulfate mimetics will alter their differentiation program by modulating Wnt signaling to produce adipocytes with increased glucose uptake and utilization.

## INTRODUCTION

Type 2 diabetes (T2D) is a growing global health problem caused by excess caloric intake, reduced energy expenditure and the resulting onset of obesity.^1, 2^ The constant nutrient influx associated with a Western diet results in high frequency of elevated blood glucose levels. This hyperglycemia demands a continuous insulin secretion from pancreatic beta cells to ensure glucose uptake for energy production and storage.^3^ The continuous insulin secretion will desensitize its perception by adipocytes, where glucose is normally stored in the form of lipids.^4, 5^ This insulin resistance coincides with the onset of T2D, leading ultimately to beta cell failure.^6, 7^ T2D patients have a high risk to develop neuropathy, retinopathy, cardiovascular disease, stroke, and poor outcomes when coping with infectious disease.^8^ Overall, this greatly reduces their of quality of life and life expectancy. As a result, T2D has been a prominent focus of medical research to find effective treatments. Current treatment strategies focused on increasing insulin perception, augmenting oxidative tissue activity or decreasing excessive food consumption or nutrient absorption, have been limited by poor efficacy or detrimental side-effects as exemplified by the COVID-19 pandemic. ^8-10^ An alternative approach to increase insulin-independent glucose clearance is to capitalize on adipose tissue expandability and its functional metabolic flexibility and reprogram adipogenesis to increase basal glucose clearance capacity in adipose tissue.^11-14^

The cellular glycocalyx has been well-established to play a regulatory role during adipogenesis and in adipocyte function and can serve as a potential target for therapeutic intervention in T2D.^15-17^ Although, the functions of specific cell-surface glycans during adipogenic programing and their impact on glucose clearance capacity of terminally differentiated adipocytes are yet to be fully elucidated. For instance, an unbiased proteomic screen of human adipose tissues and plasma identified glypican 4 (GPC4), a heparan sulfate (HS) proteoglycan, as an adipokine.^15, 18^ In patients, plasma GPC4 levels correlate positively with the severity of insulin resistance, impaired glucose uptake, and high body mass index (BMI).^15, 19^ Genetic knockout or shedding of GPC4 is associated with an overall reduction in adipogenesis *in vitro*, which manifests in concurrent reduction of glucose uptake, insulin sensitivity, and lipid accumulation in adipocytes.^19^ It is not clear whether the adipokine function of GPC4 stems from the protein core or its glycosylation modification with HS glycans.^20^

HS are polysaccharide chains comprised of repeating units of *N*-acetylglucosamine (GlcNAc) and glucuronic acid (GlcA), which are enzymatically modified through *N*-deacetylation, epimerization at the uronic acid residue, and sulfation.^15^ These modifications produce domains with negatively charged residues, which are selectively recognized by HS-binding proteins.^21^ Cell surface HS has been extensively studied in the context of promoting lipoprotein uptake and metabolism.^22^ Accordingly, defects in lipid accumulation in differentiated adipocytes lacking Syndecan-1, also an HS proteoglycan (HSPG), or the HS biosynthetic enzyme, *N*-acetylglucosamine *N*-deacetylase-*N*-sulfotransferase 1 (NDST1), have been attributed to decreased endocytosis of triglyceride-rich lipoproteins.^23, 24^ Cognizant of the regulatory roles of HS in cellular signaling and differentiation,^25^ we set to investigate the possible roles of HS during adipogenic differentiation in defining the metabolic activity of terminally differentiated adipocytes.

Using a combination of genetic and chemical approaches to manipulate HS presentation and activity in the glycocalyx of pre-adipocyte cells,^26^ we uncovered the contributions of HS in regulating Wnt signaling to establish metabolic set points after differentiation. Further, glycocalyx engineering in pre-adipocytes with HS mimetics attenuated Wnt signaling and altered their differentiation program to produce adipocytes with increased glucose uptake and utilization. Chemical engineering of the glycocalyx thus opens a new therapeutic window for improving glucose clearance in T2D patients independent of their insulin sensitivity.

## RESULTS

### Heparan Sulfate Deficiency Prevents Lipid Accumulation in Mature Adipocytes

To define the impact of HS during adipogenic differentiation on establishing the metabolic setpoint of adipocytes, we utilized immortalized mouse embryonic fibroblasts (MEFs) with defective HS biosynthesis. As a model, we utilized MEFs lacking the HS biosynthetic enzyme, *N-deacetylase-N-sulfotransferase 1 (Ndst1*), which is required for the generation of sulfate domains that modulate interaction of HS with its binding proteins.^27^ We observed a significantly diminished lipid storage in adipocytes after differentiation of these *Ndst1*-deficient (*Ndst1*^*-/-*^) MEFs (**Fig.1a-b and Extended Data Fig. 1a-c**). The lack of lipid accumulation was not the result of altered growth rate (**Fig.1c**), nor a consequence of defective adipogenesis (**Fig. 1d-k**). Comparison of several adipogenic markers in *Ndst1*^*-/-*^MEFs to WT cells indicated normal adipogenesis regardless of defects in HS biosynthesis. A similar phenotype was generated in wildtype MEFs during adipogenesis upon treatment with high dose of heparin (100 µg/ml), which is a highly sulfated form of HS that serves as a competitive inhibitor for cell surface HS (**Fig. 1a-c and Extended Data Fig. 1d**). Previous studies in pre-adipocytes indicated that exogenous heparin treatment and *Ndst1* inactivation reduced lipid accumulation by interfering with HS-mediated endocytosis of triglyceride-rich lipoproteins, including very-low density lipoprotein (VLDL).^28, 29^ In contrast, *Ndst1* inactivation in MEF-derived adipocytes had no effect on VLDL binding and uptake (**Fig. 1l**). Exogenous heparin or *Ndst1* inactivation resulted in reduced binding of lipoprotein lipase (LPL), an HS-binding protein responsible for the release of free fatty acids (FFAs) from VLDL (**Fig.1m**). This protein is not expressed in MEFs according to RNAseq analysis and thus not likely contributes to lipid accumulation. Furthermore, genetic or chemical inhibition of HS did not negatively affect FFA uptake (**Fig. 1n**), suggesting collectively that altered lipid accumulation upon HS inactivation was independent of lipid uptake.

**Figure 1.**
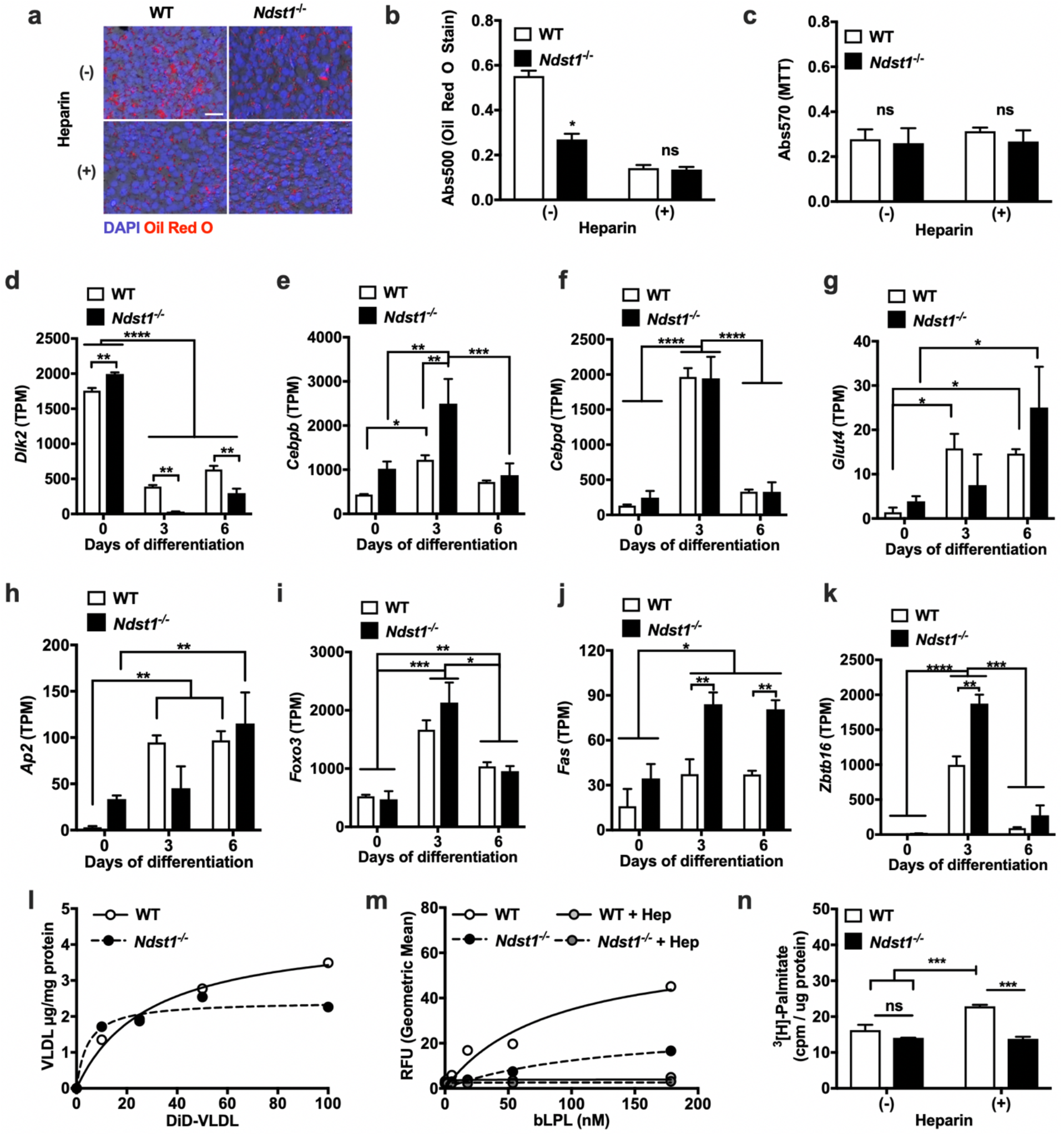
Cell surface heparan sulfate regulates lipid storage in differentiating adipocytes. **a**. Oil Red O stain of differentiated WT and Ndst1^-/-^ adipocytes treated with or without heparin (100 µg/ml). Nuclei visualized with DAPI. **b**. Quantified Oil red stain performed by elution of stain (Two-way ANOVA, n = 3). **c**, MTT assay of WT and Ndst1^-/-^ adipocytes treated with or without heparin (100 µg/ml). (n = 3). **d-k**, RNAseq quantification of adipogenesis markers relative to day 0 expression levels of WT and Ndst1^-/-^ MEFs undergoing adipogenesis (Two-way ANOVA, n = 2). **l**, Cell surface binding of Very low-density lipoprotein (VLDL) binding to the WT and *Ndst1*^*-/-*^ cells (n = 3). **m**, Lipoprotein lipase (LPL) binding to WT or *Ndst1*^*-/-*^ adipocytes treated with or without heparin (100 µg/ml) (Two-way ANOVA, n = 3). **n**, Radiolabeled fatty acid 9,10-[^3^H(N)]-Palmitate uptake in WT or *Ndst1*^*-/-*^ adipocytes treated with or without heparin (100 µg/ml) (Two-way ANOVA, n = 3). Data are presented as mean ± s.d., ^****^P < 0.0001, ^***^P < 0.001, ^**^P < 0.01, ^*^P < 0.05.

### HS Inactivation in Pre-adipocytes Alters Insulin-independent Glucose Uptake after Differentiation

Insulin-stimulated glucose clearance is one of the primary functions of adipocytes.^5^ We reasoned that the potential discrepancy between reduced lipid storage and unaltered lipid uptake could be a consequence of altered cellular glucose metabolism. To test this hypothesis, we first examined the impact of HS function in MEFs on glucose uptake in mature adipocytes. We observed a significant 84% reduction of glucose uptake in *Ndst1*-deficient adipocytes (**Fig. 2a**), which was paralleled by a 45% reduction in WT cells treated with exogenous heparin (**Fig. 2b**). Treatment of *Ndst1*^*-/-*^ with exogenous heparin did not have an additive effect on glucose uptake (**Fig. 2c and Extended Data Fig. 2**).

**Figure 2.**
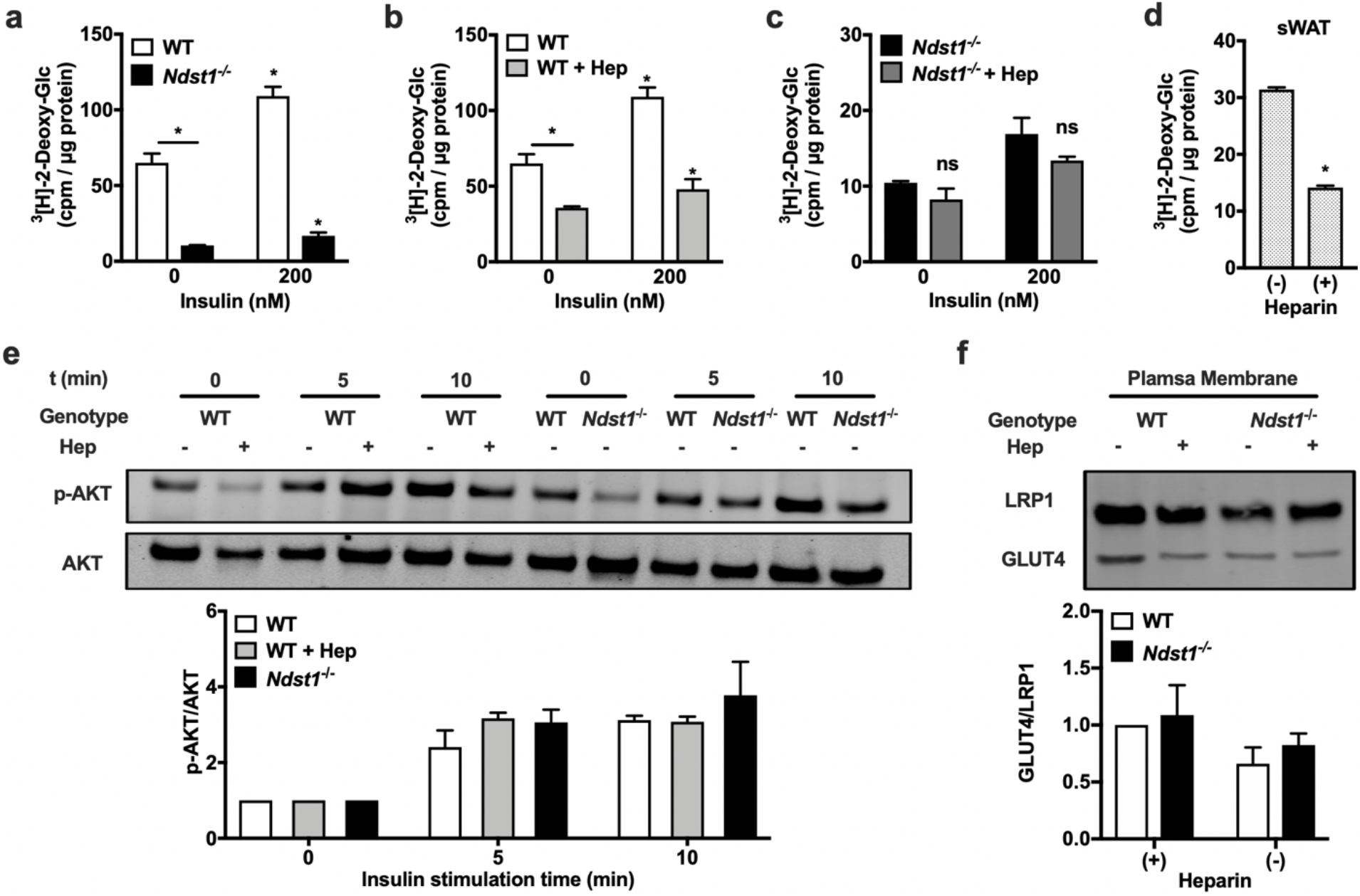
Loss of cell surface heparan sulfate interactions during adipogenesis reduce glucose uptake. **a**, *Ndst1*^*-/-*^ cells have a significantly reduced glucose uptake potential relative to WT adipocytes (Two-way ANOVA, n = 2). **b**, Heparin treatment (100 µg/mL) reduces glucose uptake potential relative to WT adipocytes (Two-way ANOVA, n = 2). **c**, Heparin treatment does not further reduce *Ndst1*^*-/-*^ adipocytes ability to access glucose (n = 2). **d**, Primary adipocytes derived from murine subcutaneous white adipose tissue (sWAT) stromal vascular cells presented with reduced glucose uptake when co-incubated with heparin during adipogenesis (Two-way ANOVA, n = 2). **e**, Stimulation of mature adipocytes (day 6) with insulin (20 nM) leads to equal signaling transduction via AKT phosphorylation regardless of HSPG interaction modulation (n = 3). **f**, Glucose transporter 4 protein (Glut4) is similarly expressed at the cell surface of WT and HSPG inhibited adipocytes (n = 3). Data are presented as mean ± s.d., ^*^P < 0.05.

We observed a similarly impaired glucose uptake in adipocytes derived from primary murine white adipose tissue pre-adipocytes differentiated in the presence of heparin (**Fig. 2d and Extended Data Fig. 3**). Surprisingly, the same differentials in glucose uptake between WT and HS-inhibited cells were maintained after insulin stimulation (−84% and - 44% respectively; **Fig. 2a-b**). This observation suggests that HS alters glucose uptake without altering insulin sensitivity. This was supported by similar levels of AKT phosphorylation in WT and HS-inactivated cells upon stimulation with insulin (**Fig.2e**). The observed diminished glucose uptake could be the result of reduced expression of cell surface glucose transporters, i.e. GLUT1 and GLUT4. However, western blot analysis of isolated plasma membrane fractions indicated unaltered levels of GLUT4 (**Fig. 2f**) and GLUT1 (**Extended Data Fig. 4**) in response to HS inactivation. These observations collectively indicate a link between cell surface HS levels and changes in glucose uptake in adipocytes, which is independent of glucose transporter expression and insulin activity.

### HS Inactivation Induces a Switch from Glycolysis to Oxidative Metabolism

A decrease in glucose uptake without changes in glucose transporter expression in HS-inactivated adipocytes can stem from reduced cellular glucose utilization.^5^ The attenuated demand for cellular glucose could be the result of a switch from glycolysis to fatty acid dependent oxidative metabolism. We suspected such a metabolic switch based on the observed reduction in lipid storage (**Fig. 1a-b**) and lactate production (**Fig. 3a-b**) in HS-inactivated adipocytes. This decreased lactate production was also mirrored by high glucose levels remaining in the culture medium (**Fig. 3c**). Typically, reliance on oxidative metabolism will drive breakdown of intracellular lipids that can be compensated for by increased fatty acid uptake from external sources. Accordingly, we observed a significant reduction in intracellular palmitate levels (**Fig. 3d**) and increased influx of isotopically labeled [U-^13^C_16_]palmitate in HS-inactivated cells (**Fig. 3e**). Under oxidative metabolism, palmitate is converted into citrate via the citric acid (TCA) cycle. In agreement with enhanced oxidative metabolism in HS-inactivated cells we observed a significant increase in isotopically labeled citrate production in our experiment (**Fig. 3f**). Collectively, these observations support a switch in the metabolic program of adipocytes induced by HS.

**Figure 3.**
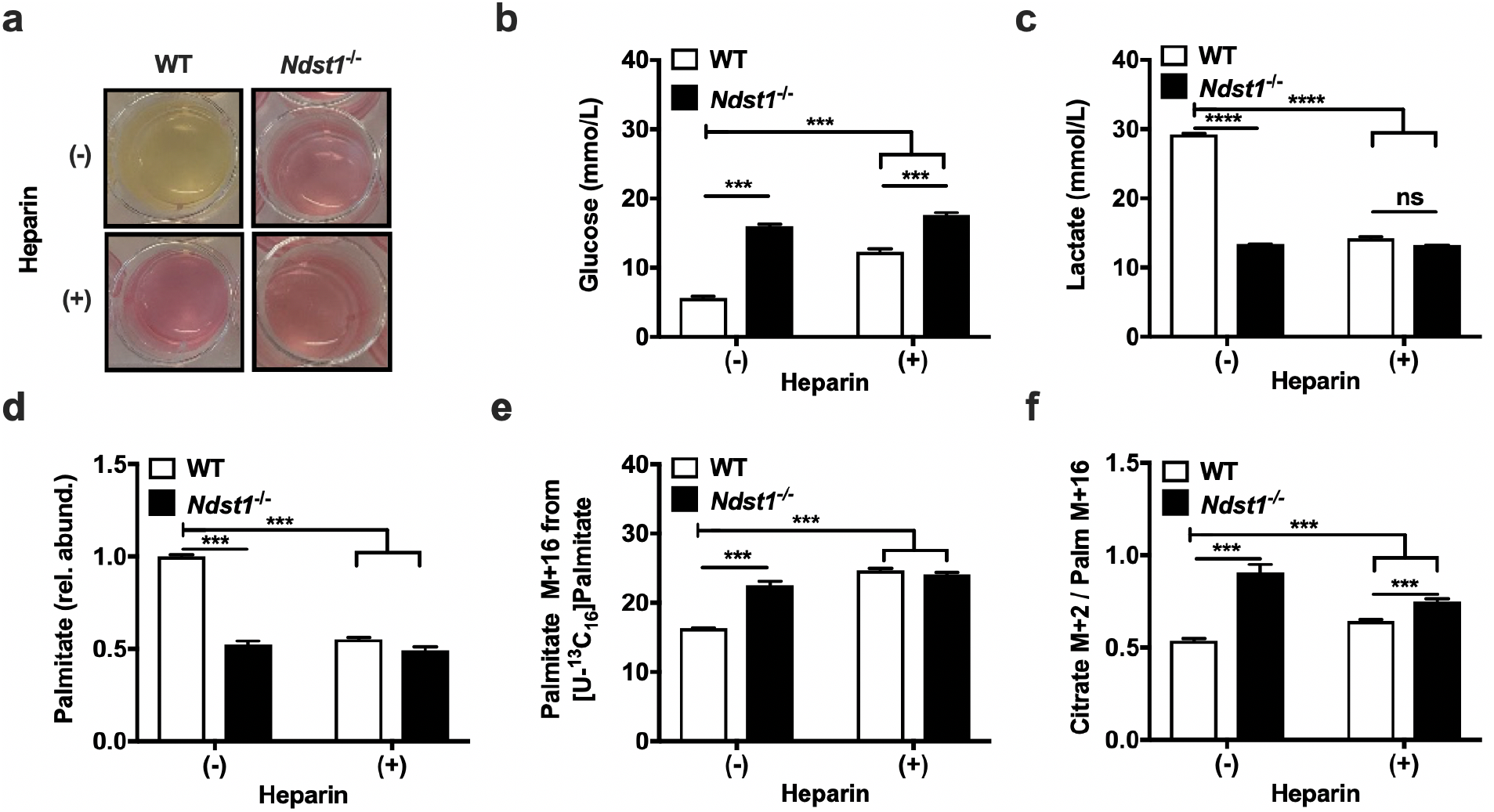
Heparan sulfate deficiency during adipogenesis promotes fatty acid oxidation. **a**, WT cells produce a greater amount of lactate than *Ndst1*^*-/-*^ adipocytes or adipocytes treated with heparin (100 µg/ml) during differentiation, as observed by discoloration of the phenol red indicator. **b**, YSI measurement of glucose in spent media collected on the final day (day 6) of adipogenesis, with fresh media at a concentration of 19 mmol/L glucose (Two-way ANOVA, n = 2). **c**, YSI measurement of lactate in spent media collected on the final day (day 6) of adipogenesis. Fresh media has a concentration of 1.5 mmol/L lactate (Two-way ANOVA, n = 2). **d**, Palmitate abundance relative to control WT cells from saponified lipids measured using mass spectroscopy, indicating total intracellular palmitate is higher in WT control cells (Two-way ANOVA, n = 3). **e**, Intracellular [U-^13^C_16_] palmitate levels indicate that HSPG-inhibited conditions present with increased palmitate uptake compared to control WT adipocytes (Two-way ANOVA, n = 3). **f**, Tracing of [U-^13^C_16_] palmitate metabolism indicates that palmitate is being increasingly converted to citrate in HSPG-inhibited adipocytes compared to control WT controls (Two-way ANOVA, n = 3). Data are presented as mean ± s.d., ^***^P < 0.001.

### HS Defines Metabolic Activity of Adipocytes During Early Adipogenesis

To determine if HS defines the metabolic switch during or after adipogenesis, we subjected mature WT and *Ndst1*^*-/-*^ adipocytes to a ^3^[H]2-deoxyglucose challenge in the presence of soluble heparin or after cell surface HS removal with heparin lyases. Neither method of acute HS inhibition resulted in altered glucose uptake (**Fig. 4a-b**), suggesting a role for HS during adipogenesis. Further support comes from lack of an effect of HS inhibition on glucose uptake in undifferentiated pre-adipocytes (MEF) or in terminally differentiated cells of non-adipogenic origin (Hep3B) (**Fig. 1c**).^30^ To define the time window during adipogenesis when HS exerts its effect, we added or removed heparin at different timepoints during differentiation (**Fig. 4d-e**). We observed that treatment with heparin for the first three days of differentiation was required to significantly reduce glucose uptake in mature adipocytes (**Fig. 4d-e**). The data point to the role for HS in defining the glucose uptake capacity early in the adipogenic program, focusing our search for a putative mechanism in this three-day differentiation window.

**Figure 4.**
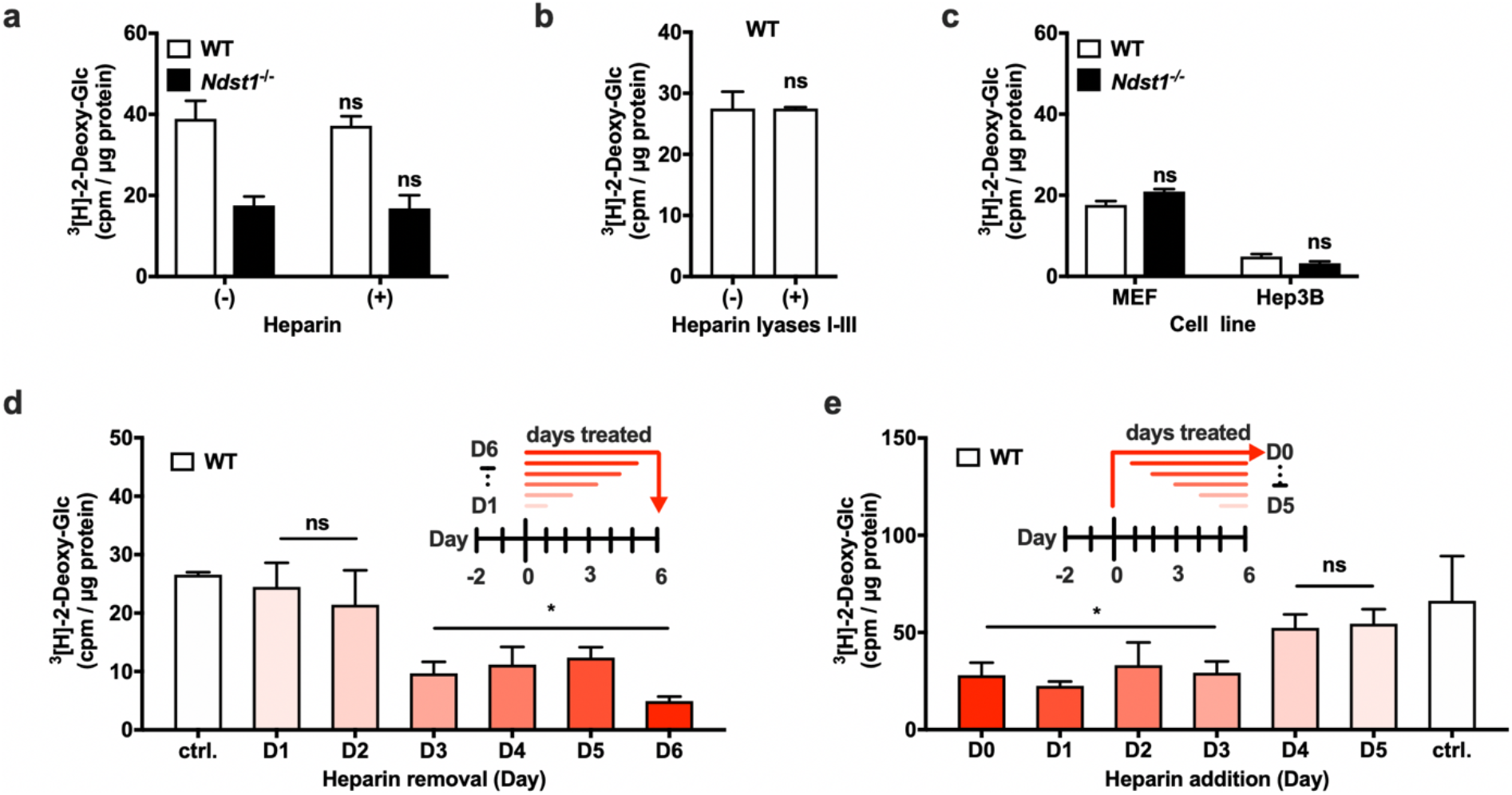
Heparan sulfate modulates adipocyte glucose clearance capacity early during adipogenesis. **a**, Adipocytes simultaneously treated with heparin (100µg/mL) during ^3^[H]-2-deoxy-glucose administration present with unaltered glucose uptake (n = 3) and independent of cell surface heparan sulfate presentation (n = 3). **b**, Treatment of adipocytes with heparin lyases I-III (30 minutes) prior to ^3^[H]-2-deoxy-glucose uptake did not altere glucose uptake (n = 3). **c**, Undifferentiated *Ndst1*^*-/-*^ MEFs and *Ndst1*^*-/-*^ Hep3B (human hepatoma) cells have no significant difference in their ^3^[H]-2-deoxy-glucose uptake capacity (n = 2). **d**, Heparin Removal: All conditions except untreated (ctrl.) cells were initially conditioned with heparin (100µg/mL) in the media. On the indicated days (day (D) 1 up to D6) heparin was removed from media to allow heparin-free differentiation conditions from that day onward. Glucose uptake potential was assessed at day 6 for all conditions. Heparin addition significantly reduced glucose uptake if cells were treated up to day 3 and beyond (Two-way ANOVA, n = 3). **e**, Heparin Addition: All conditions except untreated (day 0, D0) MEFs begin with no heparin in the media. Glucose uptake potential was assessed at day 6 for all conditions. Heparin is added at indicated days during differentiation. Addition of heparin after day 4 of differentiation can no longer impact glucose uptake in mature adipocytes (Two-way ANOVA, n = 3). Data are presented as mean ± s.d., ^*^P < 0.05 compared to control (ctrl.).

### The HS-Wnt Signaling Axis Defines the Metabolic Program in Adipocytes

HS can regulate cellular differentiation by influencing a number of signaling pathways including the FGF, SHH, TGF-ß, VEGF and Wnt superfamilies of signaling proteins.^21, 25^ To identify HS-dependent signaling pathways pertinent to metabolic programming in adipocytes, we performed comparative transcriptomic analysis of genetically (*Ndst1*^*-/-*^) and chemically (WT + heparin) HS-inhibited MEFs at early (day 3) and late (day 6) stages of adipogenesis. Overall, both genetic and chemical inactivation of HS resulted in a significant change in gene expression compared to WT controls (**Fig. 5a**). Not surprisingly, the genetic *Ndst1* deletion had a more profound overall effect on the transcriptome compared to the heparin treated MEFs. Nonetheless, both conditions resulted in a shared set of transcripts that were significantly altered compared to WT controls (**Fig. 5a**). Metascape analysis of the overlapping sets of genes indicated no major differences in typical adipogenic pathways (**Fig. 5b**).^31^ A closer examination of enriched transcripts relevant to signaling pathways regulated by HS revealed a characteristic gene signature associated with Wnt signaling (**Fig. 5b-c**).^11, 21, 25^ The genes most upregulated by HS inhibition in the early stage of adipogenesis are Wnt readout-effector genes *Plpp3, Gli3, Grk5* and *Rnf213*, indicating strong activation of the Wnt pathway (**Fig. 5c**). In agreement with increased Wnt signaling, the genes *Sox2, Lgr6* and *Klf5* were downregulated in HS inhibited cells compared to WT controls (**Fig. 5c**). These observation are consistent with the known role for canonical Wnt signaling in regulating adipocyte metabolism during adipogenesis.^11^ However, how HS influences Wnt activity in this particular process is unclear and often context dependent.^32, 33^ Our data suggests that the functional role of HS is to attenuate Wnt activity during adipogenesis.^25, 34, 35^ To confirm this hypothesis we performed differentiation experiments in WT and *Ndst1*^*-/-*^ MEFs in the presence or absence of small molecule inhibitors of Wnt. We used the canonical Wnt signaling inhibitors Niclosamide and XAV-939, as well as Wnt-C59, which additionally inhibits the non-canonical arm of the signaling pathway (**Fig. 5d-i**). All three Wnt inhibitors reversed the effect of *Ndst1* deletion on glucose uptake in adipocytes (**Fig.5e,g,i and Extended Data Fig. 5**) and showed a modest effect in WT cells (**Fig. 5d,f,h**). These experiments confirm that attenuation of Wnt signaling during early stages of adipogenesis results in enhanced glucose uptake in adipocytes after differentiation.

**Figure 5.**
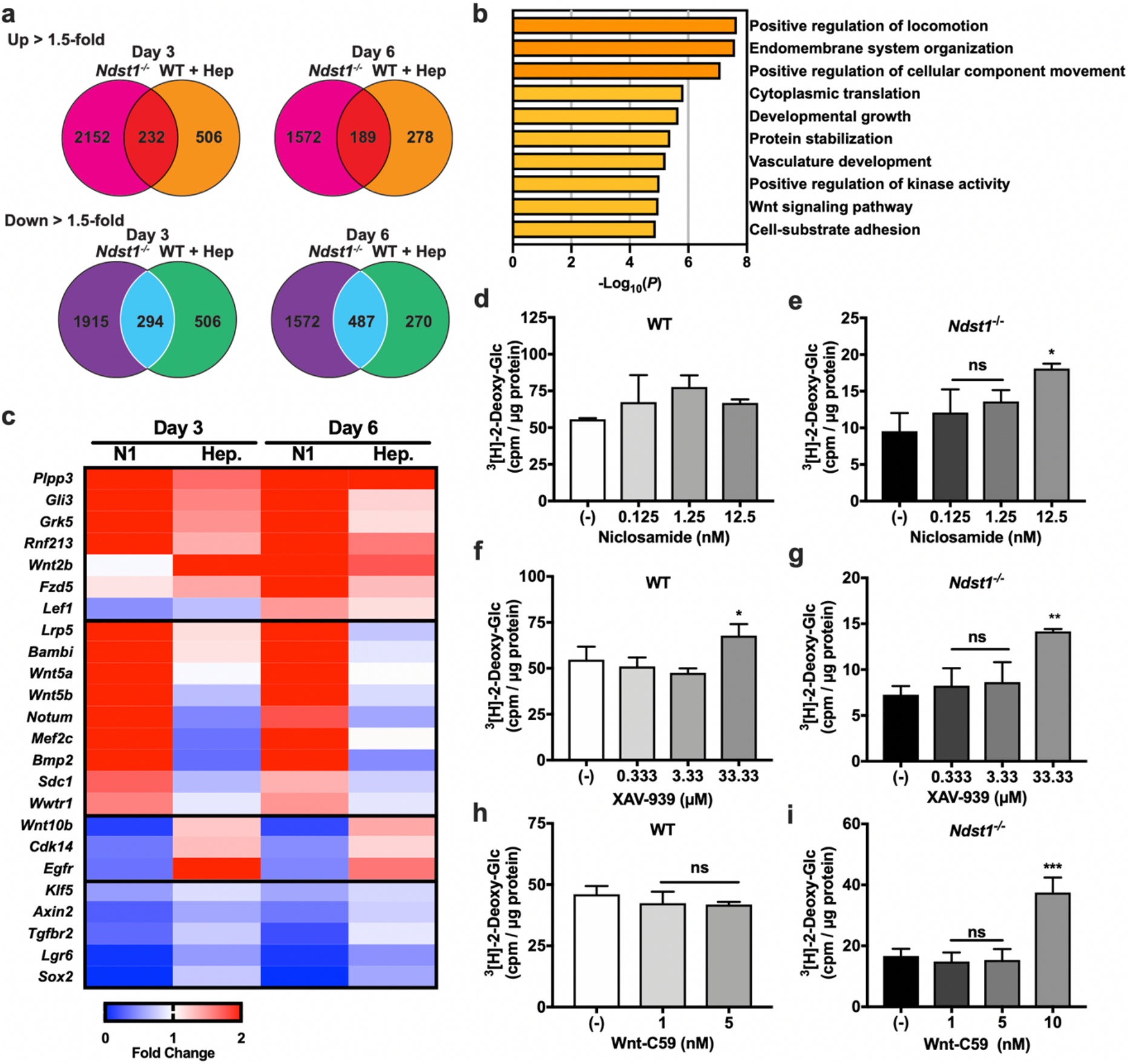
HS modulation of Wnt signaling during adipogenesis promotes glucose clearance in mature adipocytes. **a**, Venn Diagram showing the number uniquely upregulated (1.5-fold enhancement relative to WT on that day) and downregulated (1.5 fold reduction relative to WT on that day) genes on day 3 and day 6, of *Ndst1*^*-/-*^ adipocytes and WT adipocytes treated with heparin (100 µg/ml) (n = 4). Overlapping central regions indicate the number of genes up or down regulated by both *Ndst1*^*-/-*^ and WT cells treated with heparin. **b**, Metascape gene ontology analysis of genes equally affected by *NDST1* inactivation or heparin addition during adipogenesis indicate enrichment of Wnt signaling regulated genes. **c**, Heatmap of RNASeq data of Wnt related genes showing fold change in relative to untreated WT MEFs undergoing adipogenesis. **d-e**, Niclosamide, a Wnt inhibitor, treatment during the first 3 days of adipogenesis is unable to significantly enhance glucose clearance in differentiated WT adipocytes (n = 2-3). Niclosamide is able to enhance glucose clearance in *Ndst1*^*-/-*^ adipocytes at 12.5 nM (One-way ANOVA, n = 2-3). **f-g**, Treatment during the first 3 days of adipogenesis of wild type and *Ndst1*^*-/-*^ adipocytes with Wnt inhibitor, XAV-939, enhances glucose clearance in differentiated adipocytes (One-way ANOVA, n = 3). **h-i**, WT cells differentiated in the presence of Wnt-C59 had no effect on cellular glucose clearance (10 nM was toxic to WT adipocytes). Treatment of *Ndst1*^*-/-*^ cells with Wnt-C59 enhanced glucose clearance capacity in differentiated *Ndst1*^*-/-*^ adipocytes (One-way ANOVA, n = 2-3). Data are presented as mean ± s.d., ^***^P < 0.001, ^**^P < 0.01, ^*^P < 0.05.

### Chemical remodeling of the preadipocyte glycocalyx enhances glucose uptake after differentiation

Our results point to the role of cell surface HS in inhibiting Wnt activity, presumably by sequestering the ligand away from its receptor Frizzled (FZD) (**Fig. 6a**).^33^ Targeting the Wnt signaling pathway has been suggested previously as a therapeutic opportunity for increasing glucose uptake capacity in adipose tissues.^11, 36^ However, in our experiments we were able to observe only marginal benefits (up to 40% increase in glucose uptake) in WT cells in the presence of chemical Wnt signaling inhibitors at the maximum tolerated dosage. We employed an alternative strategy to augment the natural capacity of the pre-adipocyte glycocalyx to inhibit Wnt signaling in order to enhance glucose clearance. We engineered the cell surfaces of WT and *Ndst1*^*-/-*^ MEFs to present synthetic mimetics of HS.^37^ The HS mimetics comprise a synthetic linear polymer backbone decorated with disaccharide motifs representing the basic structural units of HS (**Fig. 6a**). The disaccharide side-chains were differentially sulfated to provide a range of binding avidity for the Wnt ligands. The HS mimetics carrying the most sulfated disaccharide (D2S6) showed Wnt5b and Wnt10a binding characteristics comparable to heparin (**Fig. 6b**). Additionally, the HS mimetics were endowed with a hydrophobic lipid (DPPE) anchor, for insertion into the cell membranes, and a flurophore (AF488) for quantification. Incubation of pre-adipocytes with the HS mimetics for 1 hour at 37°C resulted in efficient and dose-dependent cell membrane incorporation (**Extended Data Fig. 6**). We observed higher incorporation levels for sulfated HS mimetics in *Ndst1*^*-/-*^ MEFs compared to WT controls (**Extended Data Fig. 6**). This can be expected based on the reduced overall negative charge of HS in the glycocalyx of *Ndst1*^*-/-*^ MEFs. The remodeled pre-adipocytes were subjected to differentiation with the polymer being re-introduced once a day over the previously identified HS activity window (Day 0-3). On day 6 of differentiation the adipocytes were assayed for glucose uptake and lactate production (**Fig. 6c-d**). The sulfated Wnt-binding HS mimetics were able to restore the reduce basal glucose clearance associated with *Ndst1* inactivation. The membrane targeting of the HS-mimetics is critical for their activity as supplementation with soluble heparin had no effect (**Fig. 6c-d**). Similarly, sulfation of the disaccharide is important for activity, as the non-sulfate control HS-mimetic D0A0 showing only a limited ability to improve glucose uptake. Importantly, unlike treatment with small molecule Wnt inhibitors, cell surface engineering with HS mimetics carrying the sulfated D2A6 and D2S6 disaccharides significantly improved basal glucose uptake capacity in WT adipocytes by 39% and 47%, respectively (**Fig. 6c**). This suggests that engineering the glycocalyx of pre-adipocytes to tune Wnt signaling sensitivity may provide a new effective approach for controlling the metabolic status of adipocytes favoring glucose clearance and utilization.

**Figure 6.**
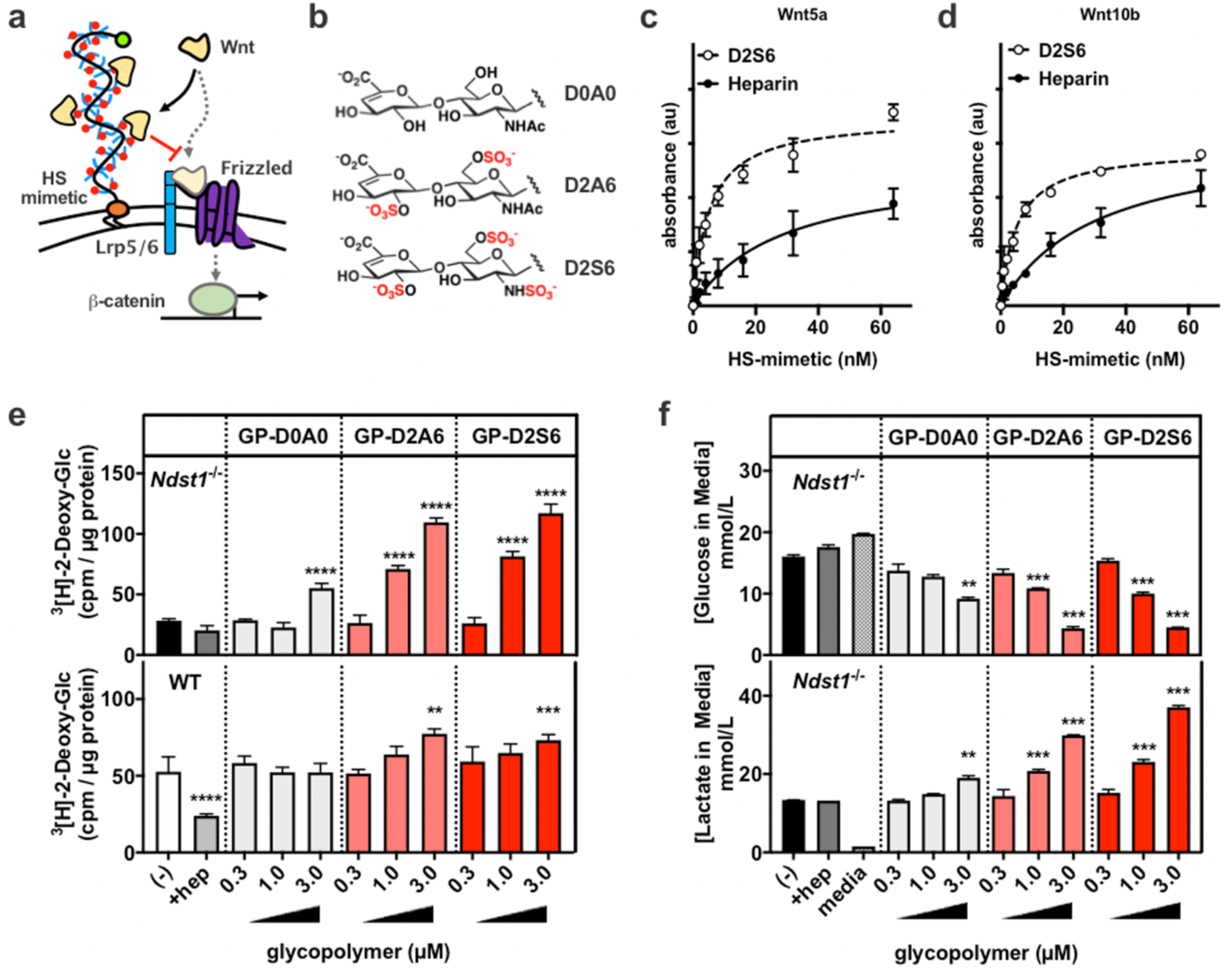
Membrane-incorporation of synthetic HS glycopolymers during adipogenesis rescues the impaired glucose uptake in HS-deficient adipocytes. **a**, Synthetic glycopolymers bearing a HS GAG disaccharide repeats were prepared and incorporated into the membranes of differentiating MEFs. Schematic of Wnt ligand sequestration by cell membrane anchored HS glycopolymers. **b**, Selected HS GAG disaccharides D2A6 (disulfated) D2S6 (trisulfated) and D0A0 (unsulfated). **c**, Wnt5a binding activity assessed by ELISA with D2S6 glycopolymer (EC_50_ = 5.1 nM, r^2^ = 0.94, n = 3) or heparin (EC_50_ = 28.2 nM, r^2^ = 0.87, n = 3), showing dose-dependent binding activity. **d**, Wnt10b binding activity assessed by ELISA with D2S6 glycopolymer (EC_50_ = 5.1 nM, r^2^ = 0.97, n = 2) or heparin (EC_50_ = 30.1 nM, r^2^ = 0.95, n = 3), showing dose-dependent binding activity. **e**, *Ndst1*^*-/-*^ and WT MEFs are treated with glycopolymer for one hour at 37 °C on day 0 to 3 of adipocyte differentiation. Sulfated glycopolymers D2A6 and D2S6 are able to dose-dependently enhance ^3^[H]-2-deoxy-glucose uptake in differentiated adipocytes (day 6) (Two-way ANOVA, n = 3). **f**, Media of treated Ndst1^-/-^ adipocytes (day 6) was assessed for glucose and lactate concentration using YSI. Cells treated with sulfated glycopolymers show a dose response rescue of glucose utilization (lowered media concentration) and increased production of the glycolysis product lactate (Two-way ANOVA, n = 2). Data are presented as mean ± s.d., ^***^P < 0.0001, ^***^P < 0.001, ^**^P < 0.01.

## DISCUSSION

A key challenge in formulating therapeutic approaches for treatment of diabetes is the management of glucose levels. The mainstream therapeutic approach is to improve insulin-mediated glucose clearance or to restrict dietary glucose intake.^9^ An alternative approach may involve reprogramming of newly generated fat tissues to increase their overall cellular metabolism and glucose demand.^11-14^ Here we present evidence that we can steer the adipogenic program to generate newly matured adipocytes with enhanced oxidative glucose metabolism to drive an increase in basal inulin-independent glucose uptake.

Using a combination of genetic and chemical tools to manipulate the HS composition on the cell surface, we identified a key role for these glycans in generating this metabolic phenotype and narrowed down the activity window during the first three days of adipogenesis. Transcriptomic analysis revealed that Wnt signaling was most perturbed in response to HS modulation within a differentiation window. Wnt signaling is a well-established HS-dependent modulator of metabolic programming during adipogenesis, and as such has been considered as a target for therapeutic intervention.^11, 25, 32, 36^ However, Wnt targeting is challenging due to its pleiotropic effects in numerous other biological systems as well as poorly defined contribution of HS function during adipogenesis.^36^

Cell surface HS can both promote and inhibit cell signaling events via either promoting signaling complex formation or by sequestering ligands away from cognate receptors, respectively.^22, 25^ In this study, we observed that eliminating cell surface HS activity pre-adipocytes led to enhanced Wnt signaling, pointing to a inhibitory role of endogenous cell surface HS in Wnt regulation. This conclusion was further supported by the restoration of the wildtype phenotype, upon application of small molecule inhibitors of Wnt signaling. Recent work identified that the protein backbone of a subclass of glypicans, including GPC4, can bind the lipid moiety of palmitoylated Wnt; serving as a ligand sequestering depot before being handed over to cognate receptors.^38^ Mutual binding of the lipid moiety to the GPC4 core protein and the interaction of protein moiety with GPC4 HS chains will increase its Wnt binding affinity and promote sequestration. However, whether such a dual binding mechanism can occur and support the central role of GPC4 as a modulator of adipogenesis remains to be established.

Although chemical inhibition of Wnt activity during adipogenesis moderately enhanced the metabolic phenotype of WT adipocytes, it required inhibitor concentrations close to the maximum tolerated dose. To overcome these limitations, we instead augmented the cellular glycocalyx with synthetic HS-mimetics with Wnt sequestering activity. The materials, based on synthetic poly(acrylamide) chains, had a Wnt binding domain composed of HS disaccharides carrying *N-*, 2-*O* and 6-*O* sulfation and a DPPE lipid for membrane targeting. This approach delivered a more robust enhancement of glucose uptake in both HS-deficient and WT adipocytes after differentiation. Interestingly, in *Ndst1*-deficient pre-adipocytes the polymer treatment resulted in a 100% improved glucose uptake capacity above the basal levels in untreated WT adipocytes. This indicates a comparable or better ability of these HS-mimetics to attenuate HS mediated signaling compared to endogenous HS in WT cells. In WT cells we observed a somewhat lower but still robust improvement in glucose uptake by approximately 50%. This outcome is consistent with less efficient incorporation of the sulfated HS-mimetics presumably due to increased electrostatic shielding from endogenous HS structures in WT cells. Further investigation should focus on optimizing the HS-mimetic architecture to further improve Wnt inhibition as well as plasma membrane targeting and localization. Our findings also open the possibility for future efforts to probe if natural variation in HS composition on adipocytes between individuals can help explain the predisposition of at-risk patients for T2D.^39^

In conclusion, we show for the first time that HS engineering can alter adipogenesis to improve glucose metabolism and clearance in newly differentiated adipocytes. This approach may open up new therapeutic avenues for the treatment of diabetes; this is particularly relevant when insulin signaling is refractory to any other treatment modality.

## ACKNOWLEDGMENTS

We like to thank Kristen Jepsen of the Institute for Genomic Medicine for performing RNA sequencing. RNA sequencing was conducted at the IGM Genomics Center, University of California, San Diego, La Jolla, CA (MCC grant # P30CA023100).

## Author contributions

G.W.T., A.R.P., K.G., and P.L.S.M.G. conceived the project; G.W.T., A.R.P., S.C.P., C.R.G., K.W., A.M., C.M.M., K.G., and P.L.S.M.G. designed and performed the experimental work.; G.W.T., A.R.P., S.C.P., C.R.G., and N.D. performed the experimental work.; K.W. and C.M.M. supplied reagents. G.W.T., A.R.P., S.C.P., C.R.G., N.D., K.W., A.M., C.M.M., K.G., and P.L.S.M.G. analyzed and discussed the data.; G.W.T., K.G., and P.L.S.M.G., wrote the manuscript. All authors discussed the experiments and results, read, and approved the manuscript.

## Materials & Correspondence

Correspondence should be addressed to: Kamil Godula, Department of Chemistry and Biochemistry, University of California San Diego, 9500 Gilman Drive #0358, La Jolla, CA 92093-0358; Philip L.S.M. Gordts, Department of Medicine, University of California, La Jolla, CA 92093, USA; +1(858)246-0994.

## METHODS

### Adipogenic differentiation

Cells were seeded on a 24-well plate at a density of 30,000 cells/cm^2^. Cells were allowed to grow to confluence for 48 hours. At this point (Day 0), the media were removed, cells were washed with PBS, and differentiation media with or without heparin (100 µg/mL) were added. Differentiation media consisted of 0.1 µM dexamethasone, 450 µM 3-isobutyl-1-methylxanthine, 2 µM insulin, and 1 µM rosiglitazone in DMEM supplemented with 10% FBS. On Day 3 of differentiation, cells were washed with PBS and treated with insulin media, which consisted of 2 µM insulin, and 1 µM rosiglitazone in DMEM with 10% FBS. The cellular cholesterol and triglyceride content was determined after lysing cells in 0.1M NaOH. Total plasma cholesterol and plasma triglyceride levels (Sekisui Diagnostics) and protein levels (BCA protein assay) were determined using commercially available kits.

### Gene expression analysis

Total RNA from homogenized tissue and cells was isolated and purified using E.Z.N.A. HP Total RNA (Omega) or RNeasy mini (Qiagen) kits according to the manufacturers’ instructions. The quality and quantity of the total RNA was monitored and measured with NanoDrop (NanoDrop Technologies, Inc. Wilmington, DE). 5-10 ng of cDNA was used for quantitative real-time PCR with gene-specific primers (**Extended Data Table 1**) and TBP as a house keeping gene on a BioRad CFX96 Real-time PCR system (Bio Rad).

### RNA-seq library preparation

Cells were lysed in Trizol and total RNA was extracted using the Direct-zol kit (Zymo Research, CA USA). On column DNA digestion was also performed with DNAse treatment. Stranded RNA-Seq libraries were prepared from polyA enriched mRNA using the TruSeq Stranded mRNA library prep kit (Illumina). Library construction and sequencing was performed by the University of California San Diego (UCSD) Institute for Genomic Medicine. Libraries were single-end sequenced for 76 cycles on a HiSeq 4000 to a depth of 20-30 million reads.

### RNASeq Analysis

The Kallisto/Sleuth differential expression pipeline analysis was performed for each of the 16 samples. Kallisto was run for single-end read quantification, using the parameters: kmer size = 31, fragment length = 280, and sd = 25. For each of the 16 samples, kallisto quantified transcript abundance with 10 bootstraps. Normalized transcript abundances were further passed into sleuth, which were then aggregated to gene level.

### Oil Red O Assay

MEFs differentiated to day 6 into adipocytes in a 24 well plate are washed twice with PBS, then fixed in 4% paraformaldehyde for 10 minutes. The cells are then washed twice with PBS and once with MilliQ water before being then incubated for 1 minute in 60% isopropanol. Then, the oil red working solution is placed on the cells for 8 minutes. The working solution is prepared from an Oil Red O stock solution consisting of 0.5 g Oil Red O in 100 mL isopropanol which has been heated to 56 °C until the Oil Red O has dissolved. The working solution is prepared by taking 30 mL of stock solution and adding to 20 mL distilled water. The mixture must stand for at least 10 minutes then be filtered before use. After the 8 minute incubation, cells are incubated in 50% isopropanol for 1 minute, followed by another 1 minute incubation in 10% isopropanol, then MilliQ water, followed by tap water. After each of these 1 minute incubations, the fixed, stained cells are placed in MQ water and imaged. The water is removed, and the wells are allowed to dry. The stain can then be eluted in 200µL 100% isopropanol over a 10 minuted incubation period. The eluted stain is then placed into a 96 well plate, and absorbance is measured at 500 nm.

### MTT Assay

MEFs differentiated to day 6 into adipocytes in a 24 well plate, and on day 6 cells are treated with 50 µL of the Cytoselect MTT assay preformulated reagent, which is added directly to media. The cells are incubated in this mixture for 4 hours as violet precipitates form. The cells are then treated with 500µL of the supplied detergent solution for 2 hours, and wells are agitated with pipetting to enhance the dissolution of the precipitate. The detergent with dissolved precipitate is then moved to a fresh 24 well plate and absorbance is measured at 570 nm.

### Lipoprotein lipase (LPL) binding assay

Bovine LPL generously provided by Gunilla Olivecrona (Department of Biomedical Sciences, Umeå University, Umeå Sweden)^42^. Enzyme activity was determined using ^3^H radiolabeled substrate as previously described^43^. Molar ratio of biotin to LPL was determined in a HABA displacement assay (Pierce Biotin Quantitation Kit).

LPL binding assays were performed similar to as previously described^44, 45^. Cells were harvested using Accutase cell detachment solution (Millipore) and washed twice with PBS. Cells were incubated for 15 min at 37°C in serum free media in the absence or presence of 5 mU/ml each of recombinant heparin lyases I, II, and III. Treated and untreated cells were washed twice with PBS, chilled on ice for 20 min and incubated with 50 nM biotinylated LPL in 1% BSA supplemented PBS at 4°C for 1 hour. Following the incubation, cells were washed twice in ice-cold PBS and incubated with 0.4 µg/mL Phycoerythrin-Cy5 conjugated Streptavidin in 1% BSA supplemented PBS at 4°C. A set of cells was exposed only to Phycoerythrin-Cy5 conjugated Streptavidin and was used as background control. Cells were washed twice with PBS and analyzed by flow cytometry.

### VLDL binding to adipocytes

Human VLDL (δ < 1.006 g/ml) was isolated from plasma by buoyant density ultracentrifugation and quantified by BCA protein assay (Pierce) as described.^30^ To label the particles, 1–2 mg of VLDL were combined with 100 μL of 3 mg/mL 1,1′-dioctadecyl-3,3,3′,3′-tetramethylindodicarbocyanine perchlorate (DiD; Invitrogen) in DMSO and then re-isolated by ultracentrifugation. After incubation with VLDL for 30 min at 4°C, cells were rinsed with PBS and lysed by adding 0.1 M NaOH plus 0.1% SDS for 40 min at room temperature.^30^ Fluorescence intensity was measured with appropriate excitation and emission filters in a plate reader (TECAN GENios Pro, Switzerland) and normalized to total cell protein.

### Glucose and Lactate Quantification

MEFs are differentiated to day 6 into adipocytes in a 24 well plate, and on day 6 500µL of media is collected from each well in an Eppendorf tube and immediately frozen in liquid nitrogen. Once all samples are collected, the frozen samples are thawed on ice and filtered through an Amicon ultra 3000 molecular weight cut-off centrifugal filter at 4 °C. Then, the filtrate is analyzed using a YSI 2900, where the sample is loaded into two wells of a 96 well plate, 200µL media per well. The samples are then analyzed for lactate and glucose concentration.

### ^3^[H]-2-Deoxy-Glucose Uptake Assay

Cells are washed twice with PBS, and placed into DMEM with 2 mg/mL BSA for two hours. Then, if cells are to be insulin stimulated, they are treated with 200 nM insulin for 30 minutes at 37 °C in freshly prepared transport solution consisting of 137 mM NaCl, 1.2 mM MgSO_4_, 1.2 mM KH_2_PO_4_, 4.7 mM KCl, 2.5 mM CaCl_2_, and 20 mM HEPES with a pH of 7.3. The insulin solution is then removed and the cells are washed twice with transport solution before radioactive transport solution is added, which consists of 0.25 µCi/well of ^3^[H]-2-deoxy-glucose and 0.025 mM 2-deoxy-glucose. After a 10 minute incubation in the radioactive transport solution, it is removed and the, the glucose uptake is halted by addition of ice cold PBS. The cold PBS is removed and each well is washed twice with PBS. Then, lysis buffer consisting of 0.1 M NaOH and 0.1% w/v sodium dodecyl sulfate, is added to each well. The radioactive lysate was transferred to a scintillation vial containing 5 mL of Ultima Gold liquid scintillation fluid and analyzed using a liquid Scintillation counter.

### Palmitate Tracer Study

MEF cells on day 8 of adipogenesis were incubated for 6h in 100uM [U-^13^C_16_]Palmitate-containing media. The media contained 10% delipidated FBS and 5% (v/v) of a BSA-conjugated [U-^13^C_16_]Palmitate stock. To make the stock, briefly, 100mM [U-^13^C_16_]Palmitic acid was dissolved in ethanol at 50°C. A solution of 4.4% essentially FA-free BSA (Sigma) in PBS was warmed to 37°C. A 50:1 mixture BSA:Palmitate solution was made and incubated at 37°C for 1-2 hours before aliquoting and freezing in glass tubes. The stock contains a 3:1 FA:BSA ratio. After incubation, cells are lysed on ice and prepared for GC-MS analysis, as previously reported.^46^

### 9,10-^3^H(N)]-Palmitate Uptake Assay

Cells are washed twice with PBS, and placed into DMEM with 2 mg/mL BSA for two hours. During this time, 14C-palmitic acid is complexed with BSA at a molar ratio of 1:1 for 30 minutes. After the cells have been starved for 2 hours, the cells washed twice with PBS and incubated in radioactive transport solution (see 2-deoxyglucose uptake assay for contents of transport solution) with 1 µCi/well ^14^C-palmitic acid:BSA for 10 minutes. After a 10 minute incubation in the radioactive transport solution, it is removed and the glucose uptake is halted by addition of ice cold PBS. The cold PBS is removed and each well is washed twice with PBS. Then, lysis buffer consisting of 0.1 M NaOH and 0.1% w/v sodium dodecyl sulfate, is added to each well. The radioactive lysate was transferred to a scintillation vial containing 5 mL of Ultima Gold liquid scintillation fluid and analyzed using a liquid Scintillation counter.

### Insulin Stimulation

MEFs differentiated to day 6 into adipocytes in a 24 well plate in the presence or absence of exogenous heparin (100 µg/mL), and on day 6 cells are washed with PBS and serum starved for 2 hours in DMEM with 2 mg/mL BSA. After 2 hours, each well is stimulated with insulin at a final concentration of 10 nM, with the insulin delivered directly to the serum free media as a 10 µL aliquot. The plate in then place into an incubator at 37 °C for the indicated amount of time. After stimulation, the entire 24 well plate is placed on ice, and the wells are washed with ice cold PBS twice. Then, 100µL RIPA buffer with protease inhibitor (PIC) and phosphostop is added to each well. The cell lysate is collected into an eppendorf, placed on ice for 30 minutes, and then centrifuged at 14,000xg at 4 °C for 15 minutes. The supernatant is then transferred to a fresh tube and protein concentration is determined using a BCA assay.

### Determination of Wnt binding to HS-mimetics via ELISA

Wnt5a and Wnt10b proteins (10 nM solution in 1% (w/v) BSA/DPBS) were immobilized on 96-well tissue culture treated plates overnight at 4°C. The plates were washed twice with 0.05% (v/v) Tween-20 in DPBS and blocked with 1% (w/v) BSA/DPBS for 6 hours at room temperature. Biotinylated HS-mimetic glycopolymers **GP** or heparin were added to the wells at increasing concentrations in 1% (w/v) BSA/DPBS. After 1 hour incubation, the wells were washed three times with 0.05% Tween-20 in DPBS. Streptavidin-HRP (1:1000 in 1% BSA/DPBS) was added for 45 min at ambient temperature. The wells were again washed three times in 0.05% Tween-20 in DPBS, followed by treatment with 100 μL of TMB substrate for 2-5 min before quenching with 100 μL of 2N sulfuric acid. Absorbance (450 nm) was measured using a Molecular Devices SpectraMax plate reader.

### Synthesis and characterization of Glycopolymers

Glycopolymers were prepared as previously reported. Protected poly(acrylamide) backbone **P** terminated with DPPE-lipid and a sulfhydryl group (**Extended Data Fig. 7-8**) (M_n_= 41,562 g/mol, M_w_= 53,061 g/mol, DP= 160, and Ð= 1.28) was used as the precursor for glycopolymer assembly. HS-mimetic glycopolymers (**GPs**) were generated from precursor **P** in a one pot synthesis through sequential chain-end labeling with AF488-maleimide reporter, side chain deprotection, and glycan ligation. The efficiency (%) of polymer chain labeling with AF488 and side-chain modification with glycans for each **GP** were determined by UV-Vis and ^1^H NMR analysis (**Extended Data Fig. 9**).

### MEF surface remodeling with membrane-targeting HS-mimetics

MEFs were cultured in 12-well plates until confluent. The cells were washed with DPBS and incubated with 200 µL solution of serum free media (DMEM) with or without the HS-mimetic glycopolymers (**GP**) at indicated concentrations for 1 hour at 37 °C. After this time, the cells were washed with DPBS and dissociated from the plate using 0.25% trypsin. HS-mimetic membrane incorporation was analyzed by flow cytometry on a BD FACS Calibur instrument, with a minimum of 10,000 events collected per condition. Data were analyzed using FlowJo software and samples were gated to a polymer untreated control.

### Adipogenic differentiation of HS mimetic-engineered MEFs

Cells were differentiated for 6 days using the standard adipogenic differentiation protocol described above. Each day on Days 0-3, the cells were washed with PBS and incubated with 200µL solution of DMEM containing **GPs** at the indicated concentrations added for 1 hour at 37 °C. After this time, the media were removed, the cells were washed with PBS, and fresh differentiation media were added. On Day 6, the adipocytes were subjected to the ^3^[H]-2-deoxy-glucose uptake assay and the spent media were analyzed for glucose and lactate content as described above.

### Statistical analysis

If not otherwise stated results are mean values ± SEM of at least three independent experiments or mice or results show one representative experiment out of three. Statistical analysis was done on all available data. Statistical significance was determined using the 2-tailed student’s t-test, one-way ANOVA followed by a Bonferroni post hoc test or two-way ANOVA to compare time courses. For statistical analysis GraphPad prism 7 software was used. ^*^ (p < 0.05), ^**^ (p < 0.01), ^***^ (p < 0.001).

