## Supporting information for "Glycocalyx engineering with heparan sulfate mimetics attenuates Wnt activity during adipogenesis to promote glucose uptake and metabolism"

***Corresponding authors:**

**Running title**: Heparan sulfate programs glucose metabolism during adipogenesis.

### EXTENDED METHODS

#### Mice

Wildtype C57BL6 mice were obtained from The Jackson Laboratory. Mice were kept on a 12/12 hour light/dark cycle and were fed *ad libitum* with water and standard rodent chow or a high-fat diet (60% fat in calories, Research Diets D12492). All animals were housed and bred in vivaria approved by the Association for Assessment and Accreditation of Laboratory Animal Care located in the School of Medicine, UCSD, following standards and procedures approved by the UCSD Institutional Animal Care and Use Committee.

#### General chemistry techniques and instrumentation

All chemicals were purchased from Sigma Aldrich and used as received unless otherwise noted. Glycans were purchased from CarboSynth (San Diego, CA) and used as received. Lipid and AlexaFluor 488-modified HS-mimetic glycopolymers **GP** and their polymer precursor **P** were prepared according to published procedures.[40] Details regarding their composition and characterization are included as Supplemental Information. HS disaccharide nomenclature was used as defined by Lawrence et al.[41] Proton nuclear magnetic resonance ($^1H$ NMR) spectra were collected on a Bruker 300MHz NMR spectrometer. Spectra are reported in parts per million (ppm) on the δ scale relative to the residual solvent ($CDCl_3$ and $D_2O$) as an internal standard. Size exclusion chromatography (SEC) was performed on a Hitachi Chromaster system equipped with an RI detector and an 8µm, mixed bed, 300 x 7.5 mm cm PL aquagel-OH mixed medium column in DMF with 0.1 % LiBr at 70 °C. UV-Vis characterization of glycopolymers were recorded with a quartz cuvette using a ThermoScientific Nanodrop 2000c spectrophotometer.

**Cell Culture**

MEF cells were grown in monolayer culture in tissue culture treated T25 flasks at 37 °C, 5% $CO_2$. Cells were maintained in DMEM +glucose, +L glutamine with 10% fetal bovine serum. Cells were passaged every 3 days at a ratio of 1:10 after dissociation with 0.25% trypsin-EDTA at 37 °C, 5% $CO_2$, which was neutralized with an equal volume of growth medium. Cells were washed with PBS after the removal of old media. Cells are seeded into a 24 well plate at a density of 30,000 cells/$cm^2$. Cells are allowed to grow to confluence for 48 hours, at which point the media is switched to differentiation media (day 0).

**Adipose Tissue Stromal Vascular Cell Differentiation**

Around 1g subcutaneous WAT from 8 to 14-week old, male or female mice was dissected, washed, minced, and digested in 1mL DMEM containing 0.25 U/mL collagenase D (Sigma) at 37 °C with constant agitation at 200 rpm for 25-35 min. To stop the digestion, complete DMEM including 10% FBS and 1% P/S was added to the digestion mixture. Then the cells were filtered through a 100 μm cell strainer. After centrifugation at 200 x g for 10 min, the floating adipocyte fraction (AF) was collected for protein isolation, while the pellet containing stromal vascular fraction (SVF) was resuspended in fresh complete medium and seeded on a 10-cm cell culture dish. At a confluency of ~80%, cells were seeded on 6- and 12-well plates and grown to confluency for differentiation. Differentiation was induced by using complete DMEM/F12 media supplemented with 1 μM dexamethasone, 0.5 mM isobutylmethylxanthine (IBMX), 5 µg/mL insulin, and 1µM rosiglitazone. Three days after induction, medium was changed to complete DMEM/F12 supplemented with 5 µg/mL insulin for two days, afterwards cells were maintained in complete DMEM/F12 medium. On day 7, fully differentiated cells were used for experiments.

**Plasma Membrane isolation**

Membrane isolation was performed according to previously published procedures.[47] Briefly, Cells from 10-cm dishes were washed with ice-cold PBS, then scraped in 1 mL hypotonic lysis medium (HLM) containing 50 mM HEPES, 50 mM sucrose, 1 mM EDTA, 100 mM NaCl and 1 x PIC and were lysed using a Dounce homogenizer (~50 strokes). Lysates were centrifuged at 5000 g, 4 °C for 10 min. The supernatant was centrifuged at 100,000 g, 4 °C for 30 min. The resulting supernatant represented the cytosolic fraction; membrane pellets were resuspended in RIPA buffer with 1x PIC for Western blotting.

**Western Blot Analysis**

Cells were lysed using RIPA buffer, and protein was quantified using a BCA assay. Protein was analyzed by SDS-PAGE on 4–12% Bis-Tris gradient gels (NuPage; Invitrogen) with an equal amount of protein loading. Proteins were visualized after transfer to Immobilon-FL PVDF membrane (Millipore). Membranes were blocked with Odyssey blocking buffer (LI-COR Biosciences) for 30 min and incubated overnight at 4°C with respective antibodies. Goat, mouse, and rabbit antibodies were incubated with secondary

Odyssey IR dye antibodies (1:14,000) and visualized and analyzed with an Odyssey IR imaging system (LI-COR Biosciences).

### EXTENDED DATA TABLES

#### Extended Data Table 1. qPCR Primers

| Gene | Forward primer (5'-3') | Reverse primer (5'-3') |
|---|---|---|
| *Glut4* | CAATGGTTGGGAAGGAAAAGGGCTA | GTAGGCGCCAATGAGGAACCGTC |
| *aP2* | ACACCGAGATTTCCTTCAAACTG | CCATCTAGGGTTATGATGCTCTTCA |
| *TBP* | GAAGCTGCGGTACAATTCCAG | CCCCTTGTACCCTTCACCAAT |
| *Pparg* | GCATGGTGCCTTCGCTGA | TGGCATCTCTGTGTCAACCATG |
| *Cebpb* | TGGACAAGAACAGCAACGA | AATCTCCTAGTCCTGGCTTG |

### Extended Data Table 2. Abbreviations

| | |
|---|---|
| AF488 | AlexaFluor488 |
| Boc | bitertbutyldicarbonate |
| BSA | bovine serum albumin |
| Đ | polydispersity index |
| DMEM | Dulbecco's modified eagle medium |
| DP | degree of polymerization |
| DPBS | Dulbecco's phosphate buffered saline |
| DPPE | 1,2-dipalmitoyl-sn-glycero-3-phosphoethanolamine |
| EDTA | ethylenediaminetetraacetic acid |
| FA | fatty acid |
| FBS | fetal bovine serum |
| GAG | glycosaminoglycan |
| GP | glycopolymer |
| GPC | gel permeation chromatography |
| HEPES | 2-[4-(2-hydroxyethyl)piperazin-1-yl]ethanesulfonic acid |
| HLM | hypotonic lysis medium |
| HS | heparan sulfate |
| IR | infrared |
| MEF | mouse embryonic fibroblast |
| Mn | number average molecular weight |
| Mw | weight average molecular weight |
| MTT | 3-(4,5-Dimethylthiazol-2-yl)-2,5-Diphenyltetrazolium Bromide |
| NaOH | sodium hydroxide |
| NDST1 | N-deacetylase-N-sulfotransferase 1 |

| | |
|---|---|
| P | protected poly(acrylamide) glycopolymer precursor |
| PAGE | polyacrylamide gel electrophoresis |
| PIC | protease inhibitor cocktail |
| PVDF | polyvinylidene fluoride |
| RAFT | reversible addition framentation chain transfer |
| RIPA | radioimmunoprecipitation assay buffer |
| SDS | sodium dodecyl sulfate |
| SEC | size exclusion chromatography |
| TMB | 3,3',5,5′-Tetramethylbenzidine |
| WT | wild type |

### EXTENDED DATA FIGURES AND FIGURE LEGENDS

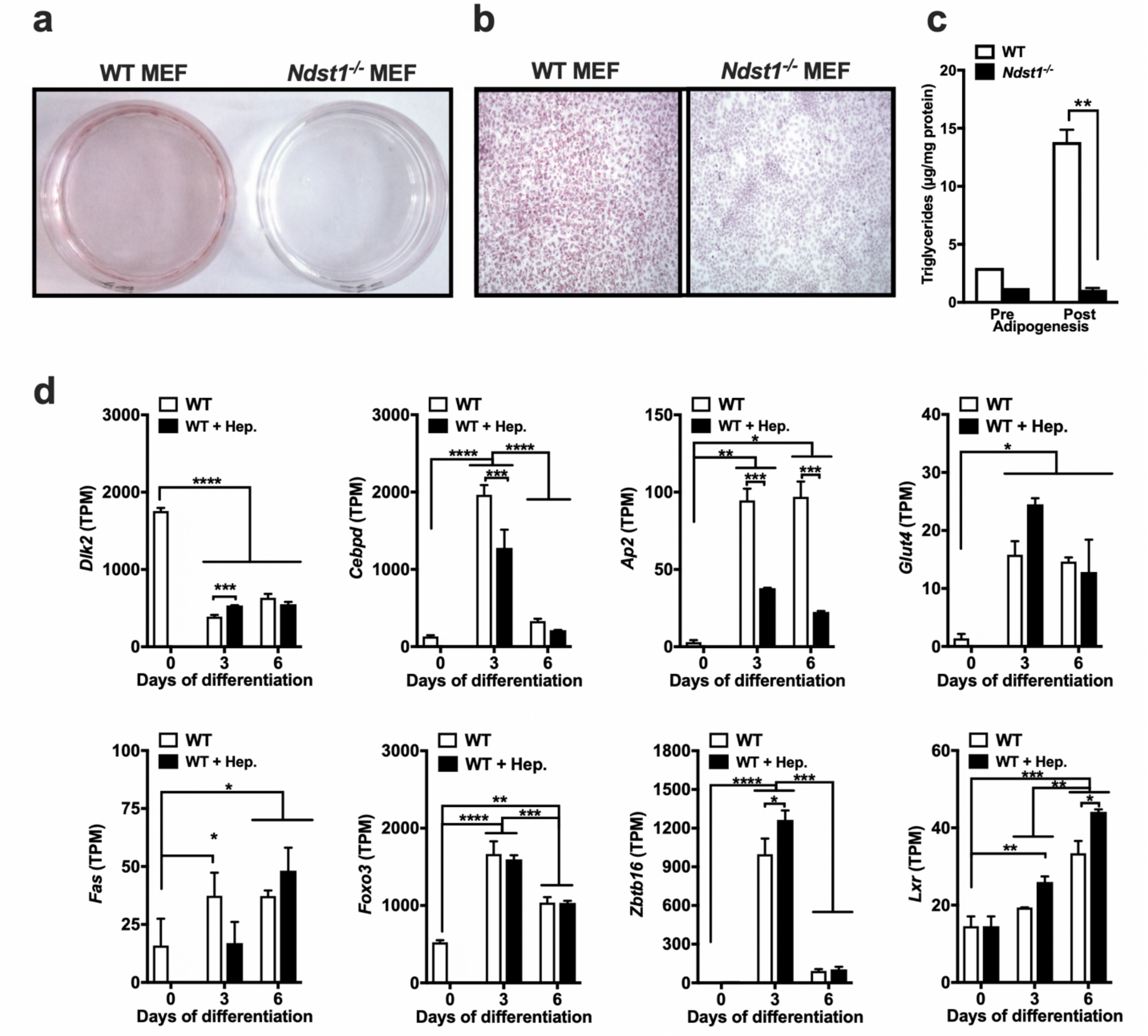


**Extended Data Figure 1. a-b.** Adipogenesis in WT and Ndst1$^{-/-}$ MEFs. Oil Red O stain and HE counter stain of differentiated WT and Ndst1$^{-/-}$ MEF. Nuclei visualized with DAPI. **c**. Analysis of triglyceride content in WT and Ndst1$^{-/-}$ MEF before and 8 days after adipogenesis (Two-way ANOVA, n=3). **d**, RNAseq analysis of adipogenic markers in differentiated WT MEFs treated with or without 100 µg/ml heparin (Two-way ANOVA, n = 2). Data are presented as mean ± s.d., ****P < 0.0001, ***P < 0.001, **P < 0.01, *P < 0.05.

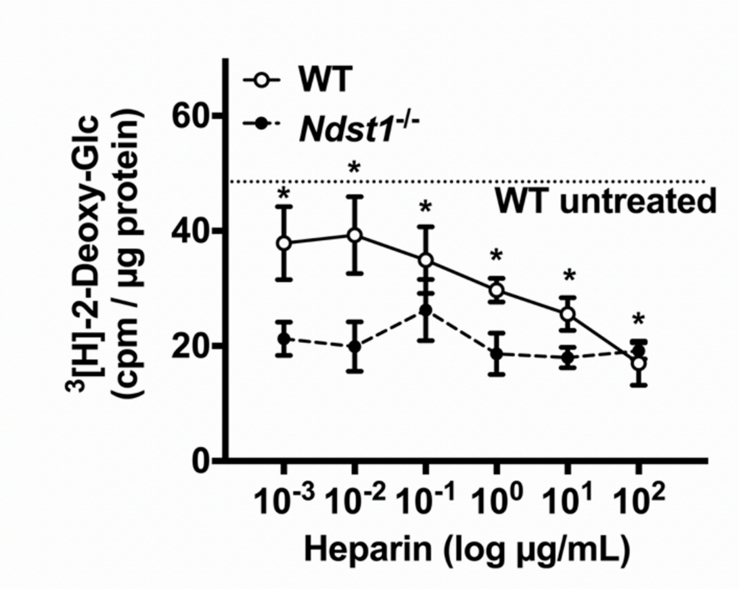


**Extended Data Figure 2.** Titration of heparin (1 ng/mL to 100 µg/mL) during adipogenesis dose-dependently reduces glucose uptake in WT MEFs, but does not affect glucose uptake in *Ndst1*$^{-/-}$ cells (Two-way ANOVA, n = 3). Data are presented as mean ± s.d., *P < 0.05.

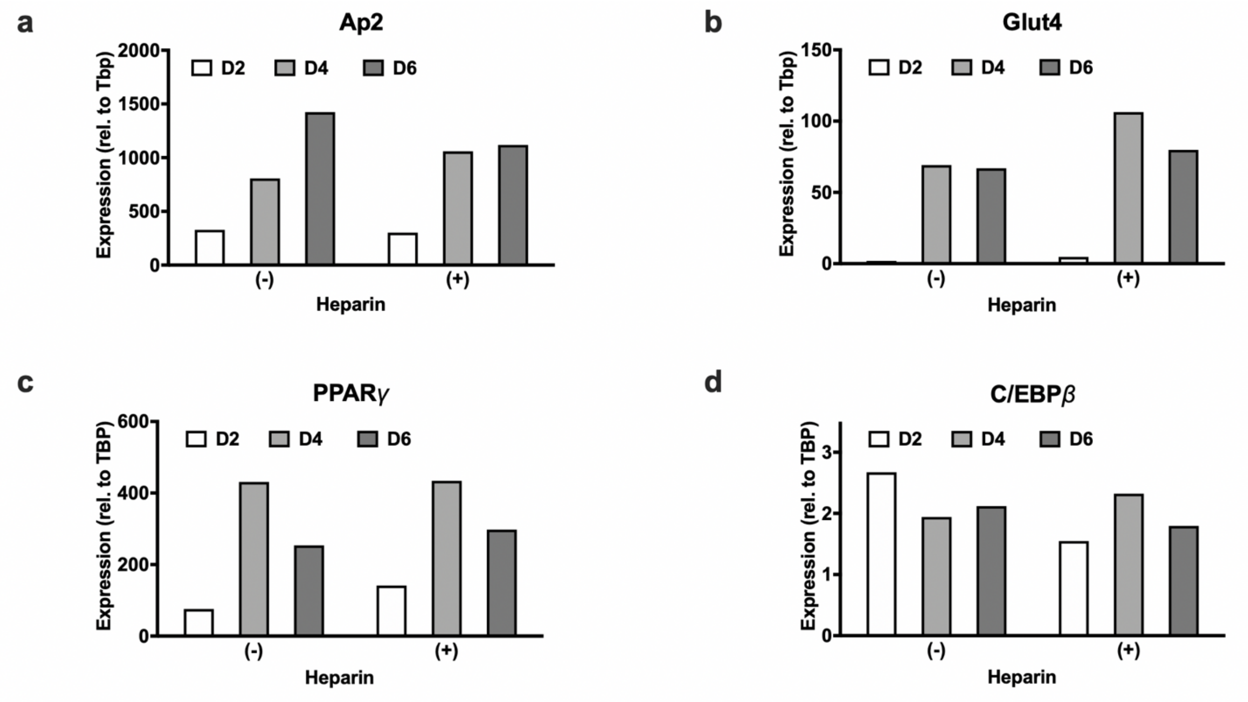


**Extended Data Figure 3.** Genetic analysis of adipogenic markers in differentiating stromal vascular cells from subcutaneous white adipose tissue at indicated times during adipogenesis (n = 2). Data are presented as mean ± s.d.

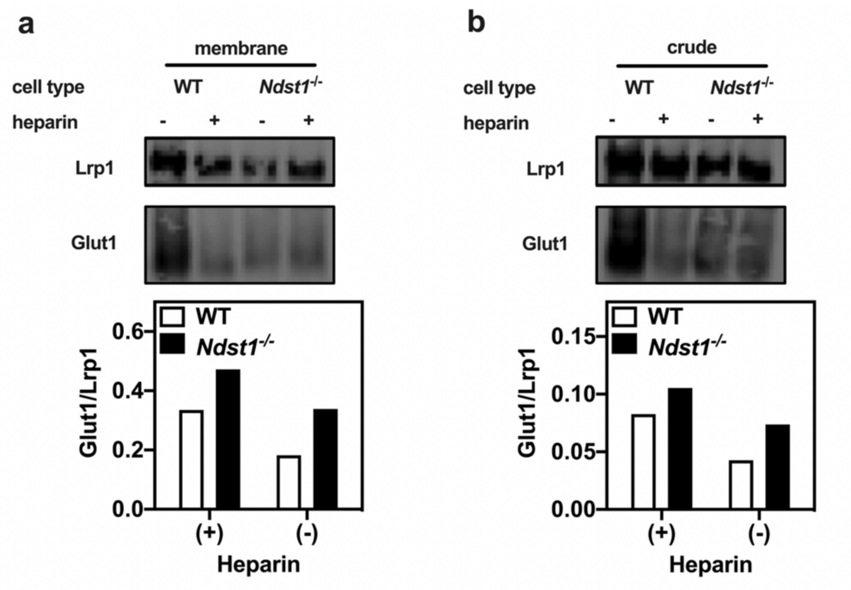


**Extended Data Figure 4.** Glucose transporter 1 (GLUT1) western blot and quantification from membrane fraction (**a**) and whole cell lysate (**b**) of day 6 WT and *Ndst1*$^{-/-}$ adipocytes. The data indicate that GLUT1 expression and membrane localization is not dramatically affected by HSPG inhibition. Interestingly, *Ndst1*$^{-/-}$ adipocytes tend to have a greater relative amount of GLUT1 compared WT adipocytes.

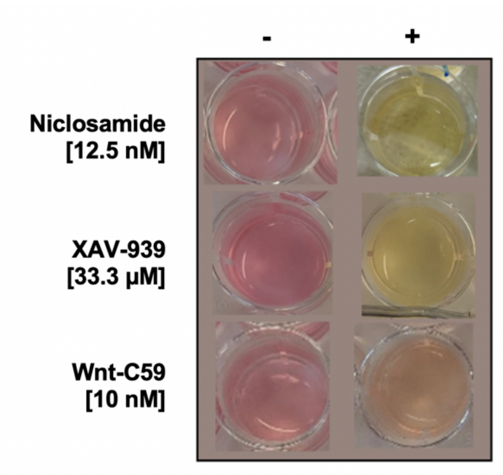


**Extended Data Figure 5.** Metabolism and Wnt signaling is affected by HSPG inhibition. *Ndst1*$^{-/-}$ adipocytes after 6 days of differentiation in the presence or absence of Wnt inhibitors Niclosamide (12.5 nM), XAV-9393 (33.3 µM) and Wnt-C59 (10 nM). Wnt inhibition (+) in *Ndst1*$^{-/-}$ adipocytes produces a greater amount of lactate compared to untreated (-) *Ndst1*$^{-/-}$ adipocytes, as observed by discoloration of the phenol red indicator.

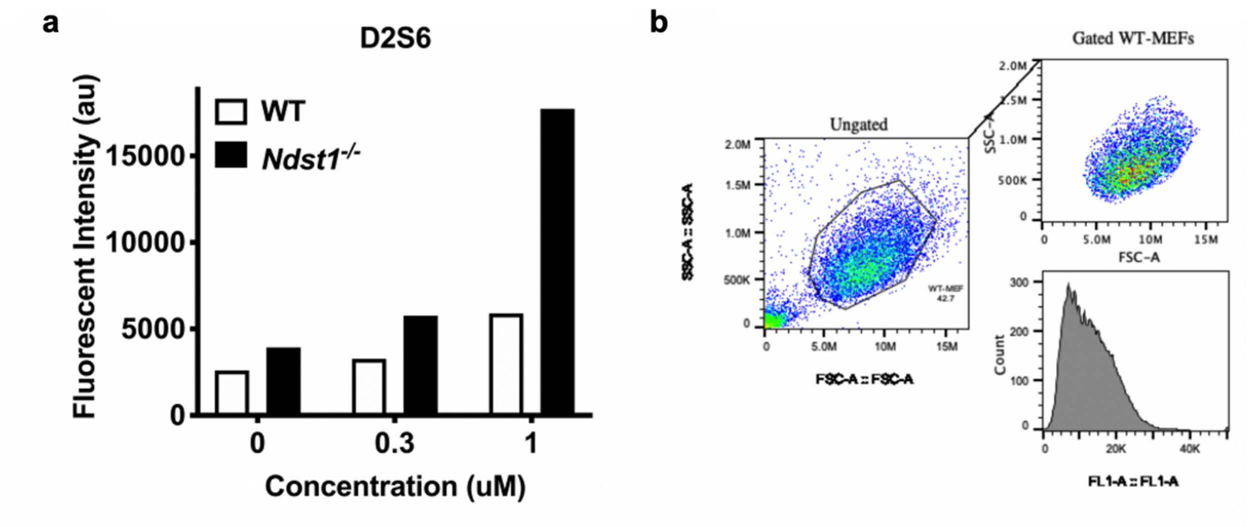


**Extended Data Figure 6.** MEF surface remodeling with membrane-targeting HS-mimetics. **a,** MEF surface remodeling with increasing concentrations of the D2S6 glycopolymer**. b**, The flow cytometry gating strategy utilized.

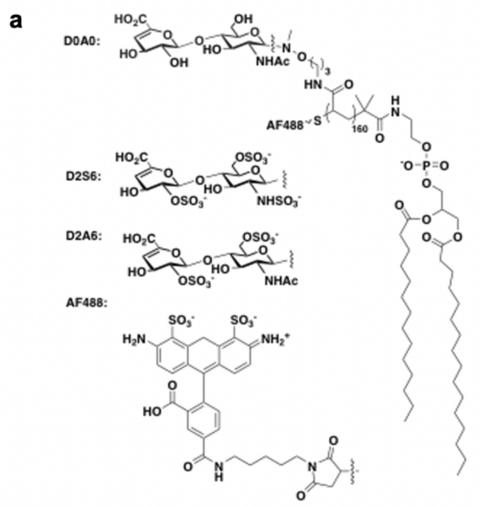


b

| GP | % AF488 | % glycan |
|---|---|---|
| **D2S6** | 86 | 43 |
| **D2A6** | 86 | 75 |
| **D0A0** | 86 | 88 |

**Extended Data Fig. 7.** HS-mimetic glycopolymer structure **(a)** and composition **(b).**

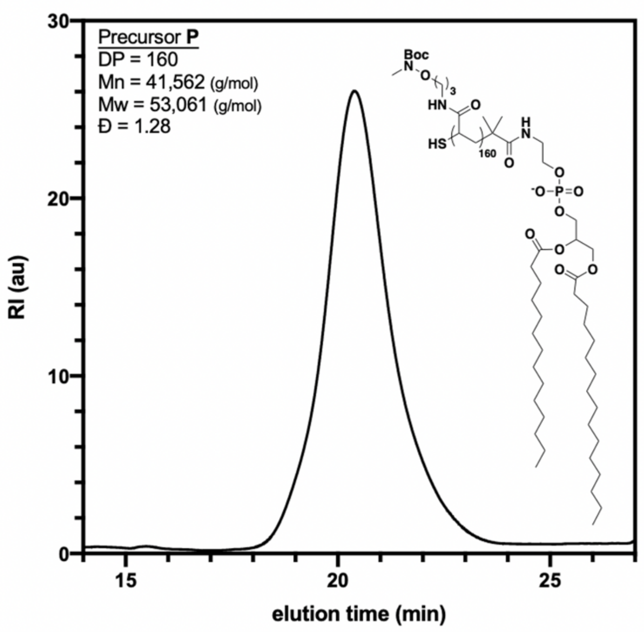


**Extended Data Fig. 8.** Structure and SEC analysis of glycopolymer precursor **P.** Precursor **P** **i**s $M_n$ = 41,562 g/mol, $M_w$ = 53,061 g/mol, DP = 160, and Đ = 1.28 as determined by SEC.

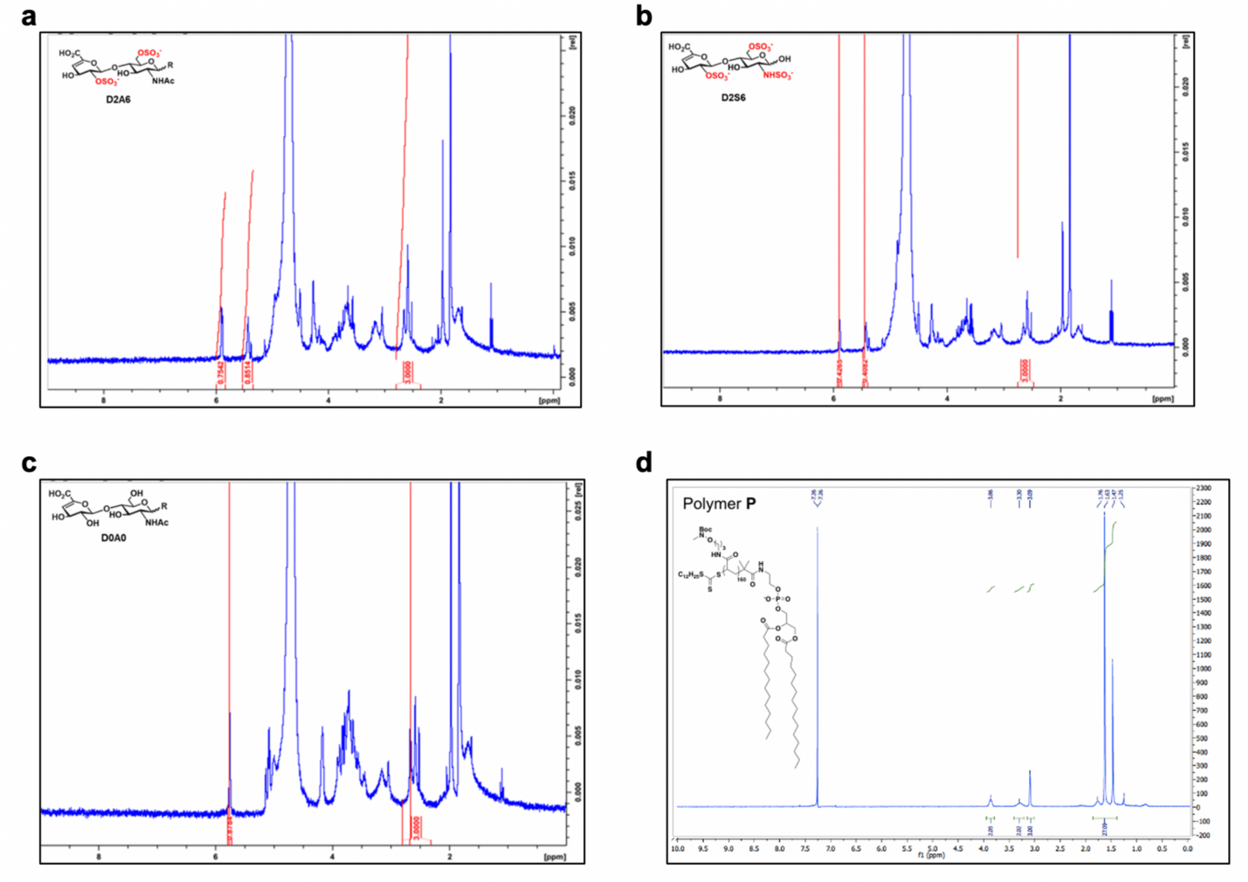


**Extended Data Fig. 9.** $^1$H NMR (300 MHz, $D_2O$) spectra for (**a-c**) HS-mimetic glycopolymers **GP** and their (**d**) precursor polymer **P**.
